# Predicting Dementia Severity by Merging Anatomical and Diffusion MRI with Deep 3D Convolutional Neural Networks

**DOI:** 10.1101/2022.08.22.504801

**Authors:** Tamoghna Chattopadhyay, Amit Singh, Neha Ann Joshy, Sophia I. Thomopoulos, Talia M. Nir, Hong Zheng, Elnaz Nourollahimoghadam, Umang Gupta, Greg Ver Steeg, Neda Jahanshad, Paul M. Thompson, the Alzheimer’s Disease Neuroimaging Initiative

## Abstract

Machine learning methods have been used for over a decade for staging and subtyping a variety of brain diseases, offering fast and objective methods to classify neurodegenerative diseases such as Alzheimer’s disease (AD). Deep learning models based on convolutional neural networks (CNNs) have also been used to infer dementia severity and predict future clinical decline. Most CNN-based deep learning models use T1-weighted brain MRI scans to identify predictive features for these tasks. In contrast, we examine the added value of diffusion-weighted MRI (dMRI) - a variant of MRI, sensitive to microstructural tissue properties - as an additional input in CNN-based models of dementia severity. dMRI is sensitive to microstructural brain abnormalities not evident on standard anatomical MRI. By training CNNs on combined anatomical and diffusion MRI, we hypothesize that we could boost performance when predicting widely-used clinical assessments of dementia severity, such as individuals’ scores on the ADAS11, ADAS13, and MMSE (mini-mental state exam) clinical scales. For benchmarking, we evaluate CNNs that use T1-weighted MRI and dMRI to estimate “brain age” - the task of predicting a person’s chronological age from their neuroimaging data. To assess which dMRI-derived maps were most beneficial, we computed DWI-derived diffusion tensor imaging (DTI) maps of mean and radial diffusivity (MD/RD), axial diffusivity (AD) and fractional anisotropy (FA) for 1198 elderly subjects (age: 74.35 +/- 7.74 yrs.; 600 F/598 M, with a distribution of 636 CN/421 MCI/141 AD) from the Alzheimer’s Disease Neuroimaging Initiative (ADNI). We tested both 2D Slice CNN and 3D CNN neural network models for the above predictive tasks. Our results suggest that for at least some deep learning architectures, diffusion-weighted MRI may enhance performance for several AD-relevant deep learning tasks relative to using T1-weighted images alone.

## 1. Introduction

### 1.1 Deep Learning in Alzheimer’s Disease Research

Alzheimer’s disease (AD) is the most common age-related neurodegenerative disease, accounting for ~70% of dementia cases worldwide. Currently, around one in three elderly people in the U.S. dies from AD or another dementia. With 6 million people now living with AD in the U.S. alone, there is an urgent need to discover factors that promote or resist dementia. We crucially need methods that detect and diagnose AD earlier and more objectively, and that facilitate disease staging and prognostic assessments. Machine learning and deep learning models have shown great promise in advancing AD research. Lam et al. [8], for example, proposed an interpretable 3D grid-attention deep neural network that can predict a person’s age and whether they have Alzheimer’s disease (AD), with promising accuracy, from their 3D structural brain MRI. This method automatically distills relevant features for classification and age prediction from large databases of raw brain MRI. In principle, this allows a direct inference of the person’s disease status without extensive image pre-processing. An MRI-based diagnostic tool could be useful in clinical practice, and in research studies, to help with screening large imaging databases and biobanks for factors in the genome or environment that might promote or resist disease. While registration of the MRI scans to a common template space, along with Gaussian mixture-based tissue classification, may further boost AD classification performance from brain MRI [2], deep learning remains attractive from a practical standpoint as it can be applied to raw images without using time-consuming parcellation and region-of-interest methods that have traditionally been used for brain morphometry. In 2020, Wen et al. [33] systematically reviewed over 30 papers applying convolutional neural networks to classify patients with Alzheimer’s disease versus matched healthy controls, based on brain imaging data. They noted the power and promise of these deep learning methods to distinguish interpretable feature patterns that are important for predicting clinical diagnosis. One remarkable recent study [2] trained a deep convolutional neural network (CNN), based on the Inception-ResNet-V2 architecture, on 85,721 brain MRI scans from 50,876 participants. After transfer learning, the model was fine-tuned for AD classification and achieved 91.3% accuracy in leave-sites-out cross-validation. In their survey, Wen et al. note that the best-performing models are not always those with the highest reported accuracy, given the pervasive use of designs that allow data leakage, and many models are not extensively tested in diverse cohorts from different populations and scanners. The stability of AD classifiers across scanning protocols and datasets has also begun to be examined. Models based on generative adversarial networks (GANs [3,4,5]) show promise in adapting data to work well with pretrained models, compensating for the so-called “domain shift [12]”.

With some recent exceptions [34,35], most CNNs for predicting AD severity have only used T1-weighted brain MRI - the most commonly collected type of brain MRI scan. Even so, we and others have recently shown [14,15] that diffusion-weighted brain MRI (dMRI), which is sensitive to subtle alterations in the brain’s microstructure, can yield metrics that are well-correlated with age, dementia severity, and even the levels of brain amyloid, a key cause of Alzheimer’s pathology which is not directly measurable on MRI. As dMRI offers independent information on white matter integrity that is not detectable with standard T1-weighted MRI, in this paper we are interested in testing what benefit dMRI might offer for AD-related deep learning tasks. Intuitively, training a CNN on diffusion MRI might perform better than training it on T1-weighted MRI alone. Also, the fusion of data from both modalities may outperform each modality used alone.

### 1.2 Introduction to Diffusion MRI and Clinical Measures of Disease Severity

Magnetic resonance imaging (MRI) is now widely used, both clinically and in research, to assist with the detection and diagnosis of a broad range of neurodegenerative conditions. In addition to T1-weighted (T1w) contrast achieved by standard anatomical MRI, other unique contrast mechanisms such as diffusion-weighted imaging (DWI) can also be used to enhance detection and improve quantification of pathology. Diffusion tensor imaging, in particular, measures characteristics of brain microstructure by modeling the local diffusion properties of water in brain tissue. While vast progress has been made in diffusion models based on multi-shell imaging protocols, Q-space imaging, and multicompartmental models of tissue biophysics [16], in this initial report we focus on the simpler diffusion metrics derived from DTI, or diffusion tensor imaging [17]. We and others have elsewhere described the limitations of DTI-derived metrics, and ways to overcome them [18]. Even so, to facilitate benchmarking of our deep learning models, here we start with the simplest metrics derived from DTI.

We briefly recap some key concepts of DTI. In the MRI scanner, the spatially-varying diffusion tensor is measured by imaging the diffusion-dependent decay of the MR signal in individual gradient directions (Fig 1). After approximating the local diffusion process by a spatially-varying diffusion tensor (essentially a 3D Gaussian approximation), there are four measures that are most commonly used to summarize the diffusion process: fractional anisotropy (FA), as well as mean, axial, and radial diffusivity (MD/AD/RD) to quantify the shape of the tensors at each voxel. These measures directly relate to the three main eigenvalues of the tensor (indicated by λ_1_, λ_2_ and λ_3_ below). These indicate the principal directions of water displacement or diffusion at each voxel. We and others have shown [15] that DTI-derived MD, RD, AD and FA maps are sensitive to aging, Alzheimer’s disease, dementia severity, and even brain amyloid burden - a key cause of Alzheimer’s disease that is not directly measurable using MRI. Fractional anisotropy is a summary measure of the directionality of diffusion, while mean diffusivity (MD) is sensitive to cellularity, edema, and necrosis. Radial diffusivity typically increases in the white matter with de- or dys-myelination. Changes in the axonal diameters or density may also influence RD. Axial Diffusivity (AD) tends to be variable in white matter changes and pathology. The standard formulae for these metrics are given below, in terms of the diffusion tensor eigenvalues.

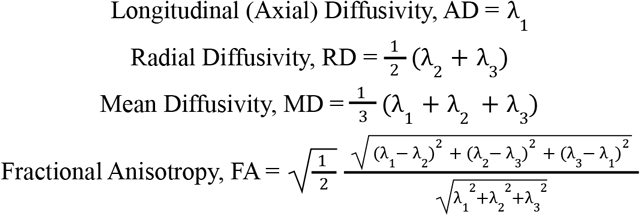

**Fig. 1.**
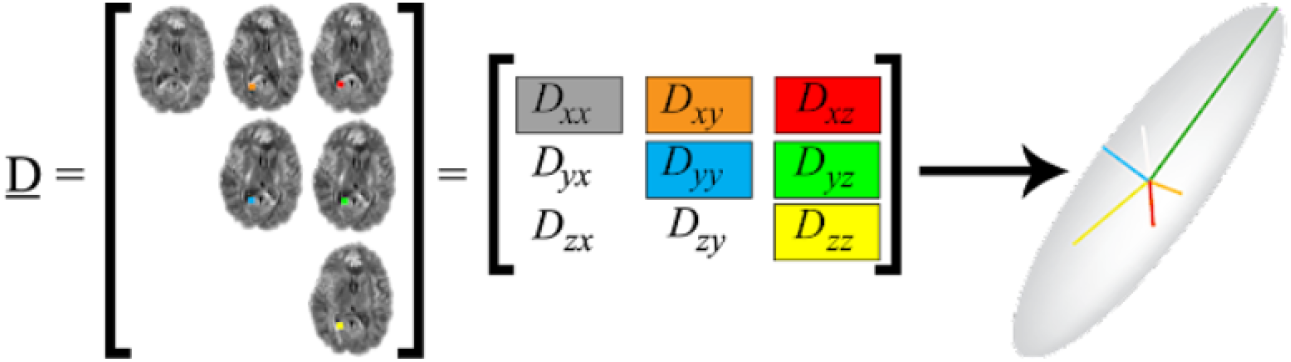
Diffusion tensor components. In the simplest case, the diffusion tensor represents a 3D Gaussian model of water diffusion, modeled using a diffusion ellipsoid, or a 3×3 matrix representing diffusion rates in different directions; this tensor can be rotated to assume a diagonal matrix form with only 3 diagonal components - the eigenvalues - which represent the relative diffusion tendencies in each of 3 principal directions.

**Fig. 2.**
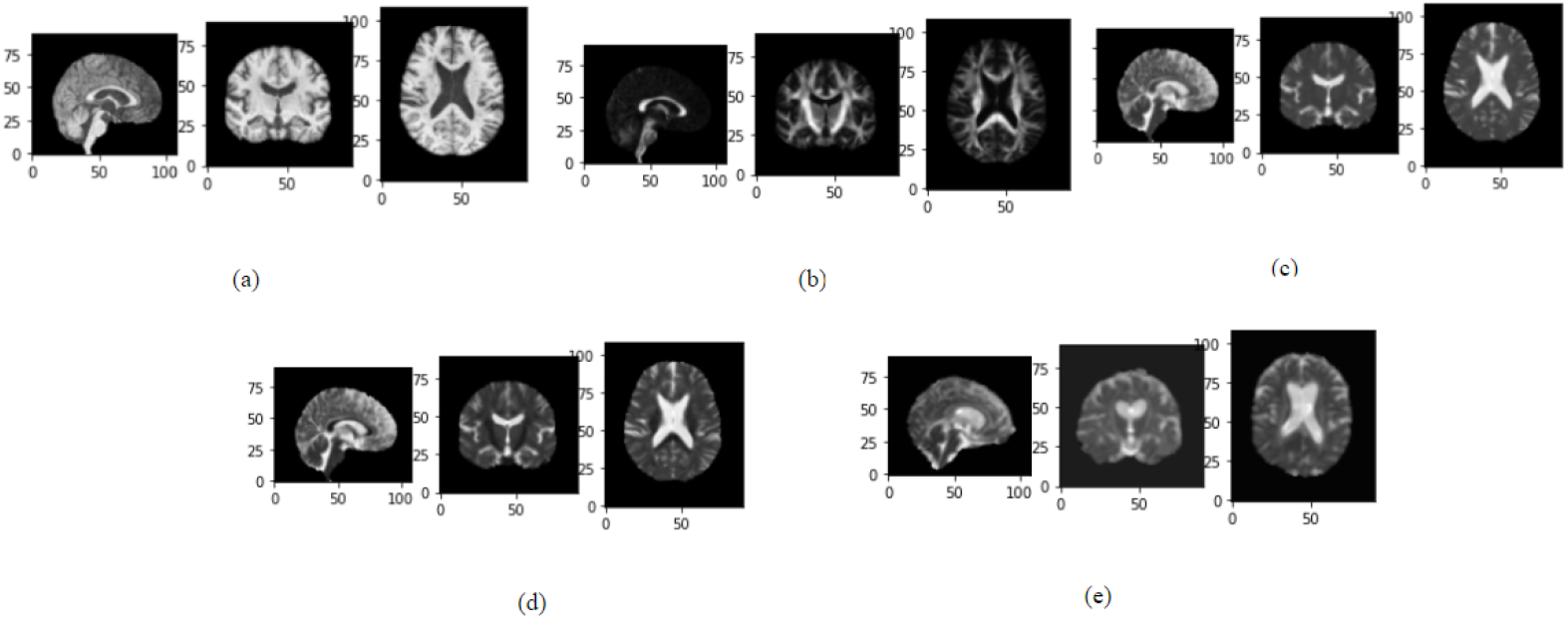
Anatomical and Diffusion MRI. Here we show example tri-axial images from (a) T1-weighted brain MRI, and DWI-derived maps of (b) FA, (c) MD, (d) RD, and (e) AD, for a typical subject from the publicly available ADNI dataset.

To benchmark our deep learning methods, we trained and tested our models on two tasks: (1) brain age prediction, and (2) dementia severity prediction. Although a person’s age is known and would not be useful to predict in clinical practice, brain age prediction in healthy control subjects is a common benchmarking task, as the ground truth (the person’s true age) is known with high accuracy. In addition, when such a trained model is applied to patients with a variety of diseases or different stages of dementia, the difference between the predicted age (the person’s “BrainAge”) and their true chronological age has been linked with future clinical decline, dementia, and mortality. This makes it a promising biomarker of brain aging, albeit with some caveats regarding pitfalls and common fallacies in interpreting this metric [19]. BrainAge is typically predicted from T1-weighted brain images, but DTI-derived metrics can also be used for this prediction task.

For predicting dementia severity, we predict Alzheimer’s Disease Assessment Scale-Cognitive Subscale test (ADAS) and Mini-Mental State Exam (MMSE) scores from the MRI scans. The Alzheimer’s Disease Assessment Scale-Cognitive Subscale test (ADAS) is one of the most frequently used tests to measure cognition in research studies and clinical trials for new drugs and other interventions. The ADAS test is more thorough than the Mini-Mental State Exam [20], and it primarily measures language and memory. ADAS11 consists of 11 parts, whereas ADAS13 consists of 13 components.

The ADAS-Cog was developed as a two-part scale, measuring cognitive and non-cognitive functions such as mood and behavior. Most current research uses the ADAS-Cog - the subscale that measures cognitive ability. ADAS-Cog 13 includes tests of attention and concentration, planning and executive function, verbal memory, nonverbal memory, praxis, delayed word recall, and number cancellation or maze tasks; the scores range from 0 to 85. The ADAS-Cog 13 is arguably more sensitive to disease progression than the ADAS-Cog 11 in subjects with AD, and similarly or slightly more sensitive in subjects with pre-dementia syndromes, such as MCI.

The Mini-Mental State Examination (MMSE) is a test used to assess cognitive decline. Patients are asked questions related to memory, concentration and ability to follow instructions. The answers are scored, with higher scores denoting relatively intact cognitive function, and lower scores signaling more severe cases of dementia. Patients with AD may decline by 2-4 points on the MMSE per year [21]. The MMSE has a maximum score of 30 points. The scores are generally grouped as follows: 25-30 points: normal cognition; 21-24 points: mild dementia; 10-20 points: moderate dementia and 9 points or lower: severe dementia.

## 2. Imaging Data and Preprocessing Steps

We use the Alzheimer’s Disease Neuroimaging Initiative (ADNI) dataset, a multisite study, launched in 2004, that aims to improve clinical trials for the prevention and treatment of Alzheimer’s disease [22]. The dataset contains subjects assessed using MRI, PET, and clinical evaluations, as well as a wide range of genetic, blood-based biomarkers, and proteomic assays. ADNI has data from a considerable number of participants with both T1-weighted MRI and DWI scans, yielding a total of 1198 subjects (age: 74.35+/-7.74 years; 600 F/598 M) with a distribution of (636 CN/421 MCI/141 AD) for our analysis.

The T1-weighted (T1w) brain MRI volumes were pre-processed using a sequence of steps, including nonparametric intensity normalization (N4 bias field correction), ‘skull stripping’ for brain extraction, nonlinear registration to a template with 6 degrees of freedom and isometric voxel resampling to 2 mm. The pre-processed images are of size 91×109×91. The T1w images were scaled to take values between 0 and 1 using a min-max scaling approach. All the T1w images were aligned to a common template.

T1w images were then nonlinearly registered to the DWI, which were subsequently warped to a common template. The diffusion MRI processing pipeline is extensively detailed in [14, 15]. Briefly, ADNI scans subjects across 58 sites with a total of seven diffusion-weighted MRI protocols, with slight customizations for sites with GE, Siemens, and Philips scanners (see **Table 1** in Zavaliangos-Petropulu et al.). The protocols vary in angular resolution, scan duration, and in the number and distribution of diffusion-weighted gradients. One example is the use of a single higher-end protocol with 127 diffusion-weighted gradients at some Siemens sites. In the current paper, we do not treat these protocols differently, but we note that it is possible to harmonize data from different protocols, using methods such as ComBat, ComBat-GAM, and CovBat. Other dMRI harmonization methods, such as VAE-GANs [23] or spherical harmonic-based models [24] will be interesting to examine in future work, as they may enhance the accuracy of CNN-based predictions from multisite dMRI.

**Table 1.** Results for brain age prediction on CN data for the 3D CNN

|  | T1 | DWI-FA | DWI-MD | DWI-RD | DWI-AD |
| --- | --- | --- | --- | --- | --- |
| Epoch = 50 \| MAE | 5.81 | **3.19** | 4.34 | 4.11 | 4.19 |
| Epoch = 75 \| MAE | **5.47** | 4.36 | **4.31** | **3.99** | **3.69** |
| Epoch =100 \| MAE | 5.63 | 3.39 | 4.72 | 4.37 | 4.13 |

**Table 2.**
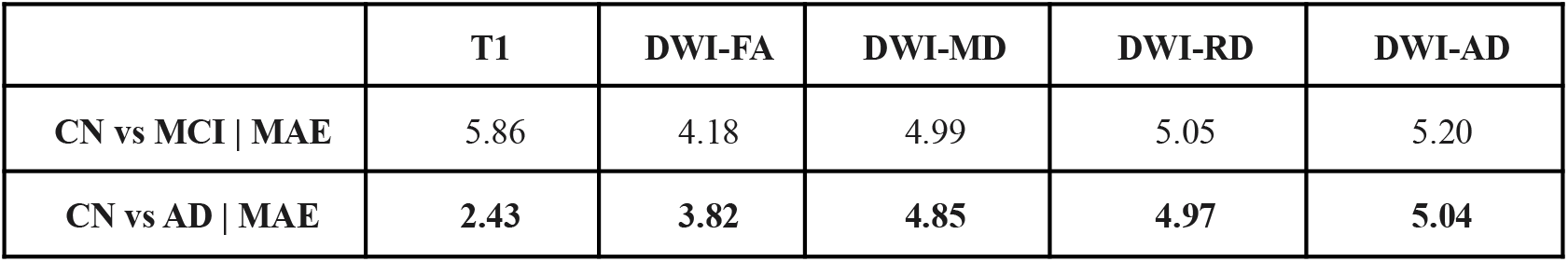
Results for linear regression to predict delta brain age gap

**Table 3.** Results for brain age prediction on CN data for the 2D slice network

|  | T1 | DWI-FA | DWI-MD | DWI-RD | DWI-AD |
| --- | --- | --- | --- | --- | --- |
| MAE | 4.89 | 7.01 | 5.01 | 4.32 | 4.55 |

**Table 4.** Results for brain age prediction on CN data for stacked dual modality using 3D CNN architecture

|  | **T1 + DWI-FA** | **T1 + DWI-MD** | **T1 + DWI-RD** | **T1 + DWI-AD** |
| --- | --- | --- | --- | --- |
| **MAE** | 6.14 | **5.44** | 5.95 | 5.62 |

**Table 5.** Results for brain age prediction on CN data for concatenated dual modality using 3D CNN architecture

|  | **T1 + DWI-FA** | **T1 + DWI-MD** | **T1 + DWI-RD** | **T1 + DWI-AD** |
| --- | --- | --- | --- | --- |
| **MAE** | 5.04 | 5.95 | 6.68 | **4.95** |

**Table 6.** Results for cognitive score prediction from T1w MRI and DWI

|  | **T1** | **DWI-FA** | **DWI-MD** | **DWI-RD** | **DWI-AD** |
| --- | --- | --- | --- | --- | --- |
| **ADAS 11 \| MAE** | 3.66 | 3.44 | 3.04 | **3.02** | 3.16 |
| **ADAS 13 \| MAE** | 5.57 | 5.01 | 4.62 | **4.58** | 4.73 |
| **MMSE \| MAE** | 2.41 | 2.38 | 1.64 | **1.54** | 1.66 |

## 3. Models and Experiments

### 3.1 3D CNN Architecture using T1w and DWI for Brain Age Prediction in Healthy Control (CN) Participants

For our first task - predicting brain age in healthy controls - we analyzed T1w and DWI images from 636 subjects (CN, mean age: 73.48+/-7.24 yrs., 375 F/ 261 M). The data was split into independent training, validation and testing sets in the ratio 70:20:10. The 3D CNN (Fig 4 top part) had three convolution blocks with filter sizes 32, 64 and 128, and additional layers of instance normalization and max pooling. We trained this CNN model for 50, 75 and 100 epochs, with a learning rate 1×10^−4^, and a batch size of 8. The learning rate is exponentially decayed with a decay rate of 0.96. The Adam optimizer [25] and mean square error loss function were used for training. To deal with overfitting, dropout between layers and early stopping were used. Test performance was assessed using mean absolute error (MAE) to compare results for different modalities.

**Fig. 3.**
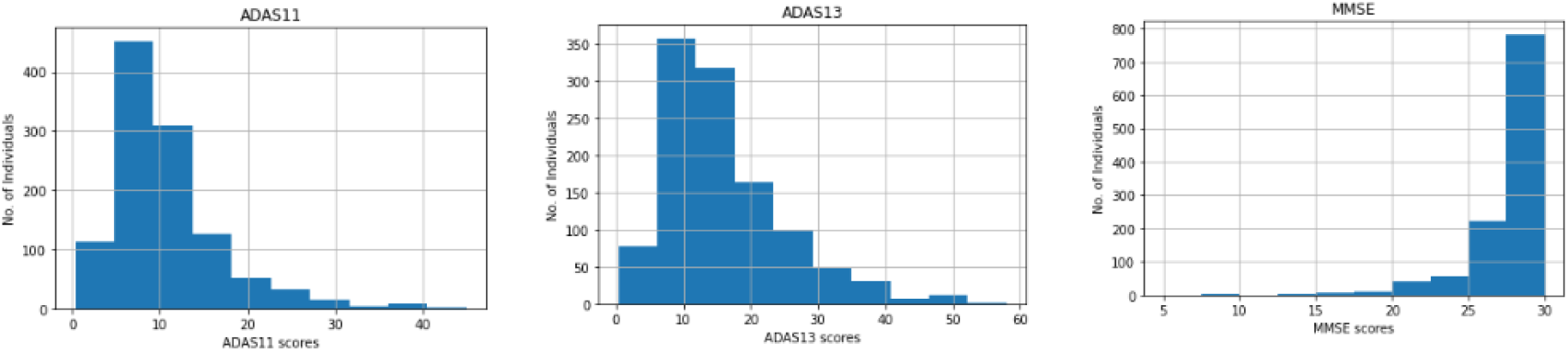
Distribution of clinical scores for the participants analyzed in this study. The ADAS scores tend to show less ceiling/floor effect than the MMSE scores, but they are still somewhat skewed (towards better performance).

**Fig. 4.**
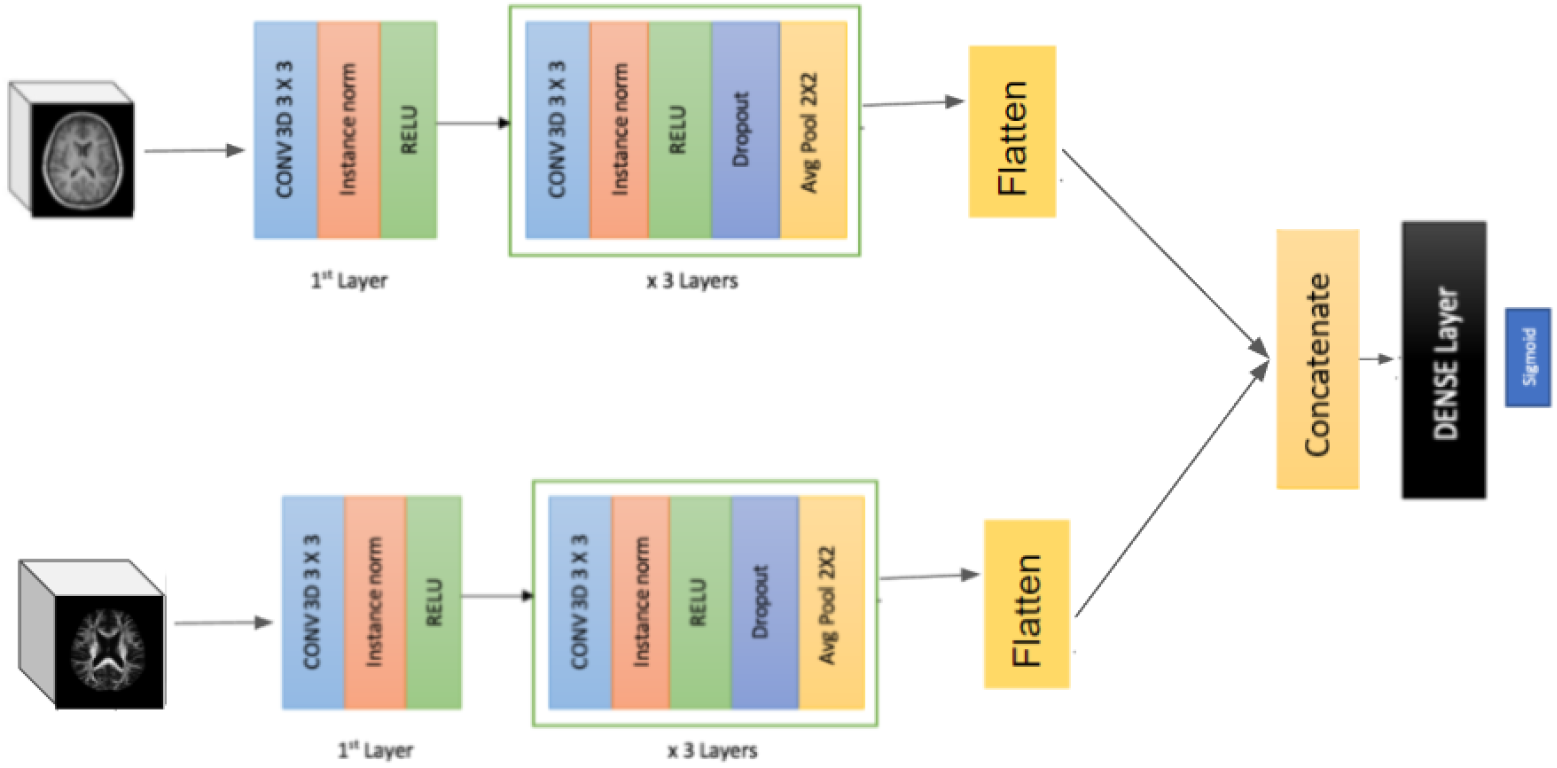
The 3D Convolutional Neural Network architecture.

**Fig. 5.**
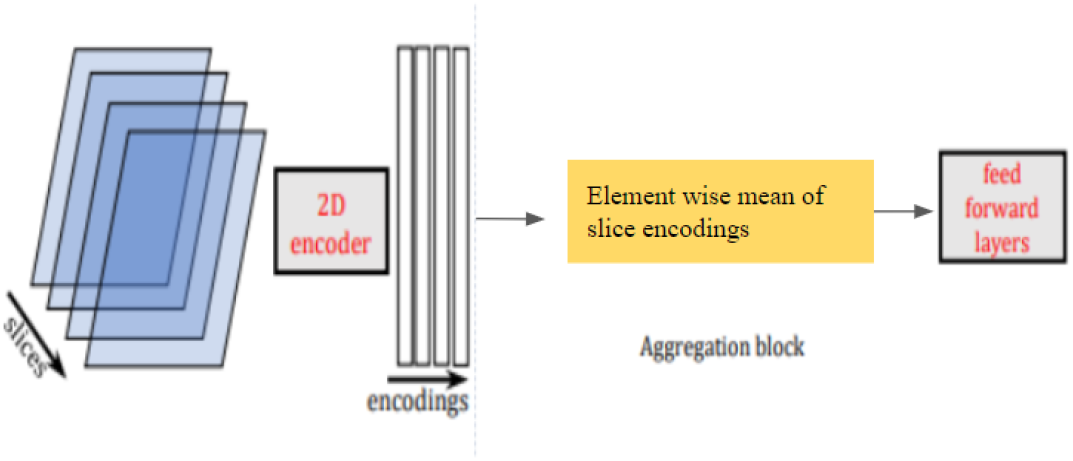
Model architecture with mean-based aggregation. Gray blocks are trainable parameters, whereas yellow blocks are operations only.

**Fig. 6.**
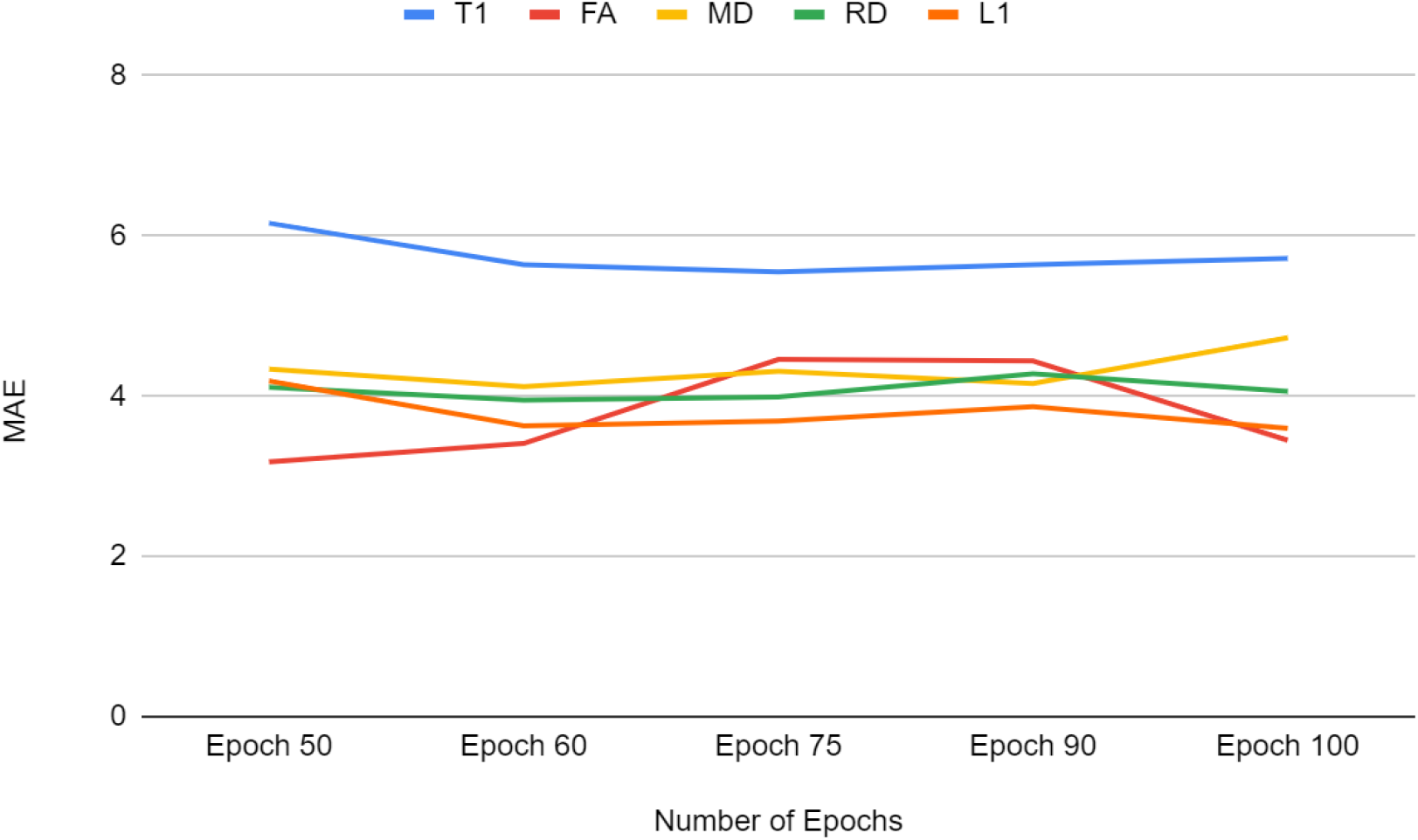
MAE vs number of Epochs for each modality.

### 3.2 2D Slice Network Architecture using T1w and DWI for Brain Age Prediction on CN data

The same CN data from experiment 3.1 was used in experiment 3 to compare fully-3D CNNs with 2D CNN networks. As noted in many prior works, a 3D image can be analyzed using a fully 3D CNN, using 3D kernels in the convolution layers. Alternatively, a 2D CNN can apply 2D kernels to multiple slices in the 3D image, and then subsequently merge the results, using methods such as set networks or permutation invariant layers [9]. The use of 2D convolutions can greatly reduce the memory requirements, the number of free parameters in the network, and training time, and can also allow pre-training on very large databases of 2D photographic images, such as ImageNet.

Here we adopted the 2D CNN by Gupta et al. [9] which was previously benchmarked for brain age estimation, but not with diffusion MRI. The model takes 3D scans as input and encodes each slice using a 2D-CNN encoder. The slices are combined using an aggregation module via permutation invariant layers resulting in a single embedding for the scan. This embedding is passed through the feed-forward layers to predict the person’s age from their MRI. The model is trained end-to-end using mean squared error loss.

The 2D CNN encoder takes a single 2D slice as input and outputs a *d*-dimensional embedding for each slice. The number of filters in the last layer of the architecture is *d* - which is decided by the dimension of the output of the aggregation module. Permutation invariant layers are used as the aggregation module, as this makes the output of the module independent of slice order. The element-wise mean of all the slice encodings was computed and used as the permutation invariant layer. *d* was kept fixed at 32, and one hidden layer network with 64 activations was used as the feed forward layer. The slices were sagittal. The model was trained for 100 epochs with the Adam optimizer, a weight decay of 10^−4^, a learning rate of 10^−4^ and a batch size of 8, with mean squared error loss.

### 3.3 3D CNN to predict Brain Age using stacked dual modality data from healthy controls

The data and 3D CNN architecture from experiment 3.1 were used to predict each subject’s age from dual modality data - i.e. the modalities were fused and used as inputs. This was done using two different ways: the T1w image data was stacked together with different DWI-derived scalar maps, and this concatenated data was fed into the 3D CNN as input, and the other way was feeding two different modalities into the CNN architecture and concatenating the results at a lower layer to run through dense layers (Fig 4). The model was trained for 50 epochs, with a learning rate 1×10^−4^, and a batch size of 8. The Adam optimizer and mean squared error loss were used during training. To deal with overfitting, dropout between layers and early stopping were used. Test performance was assessed using mean absolute error, to compare results for different modalities.

### 3.4 3D CNN to predict clinical scores from T1w and DWIs

The whole dataset of 1198 subjects. was used along with the 3D CNN architecture from experiment 3.1 to predict the clinical score values for ADAS11, ADAS13 and MMSE. The model was trained for 50 epochs, with a learning rate 1×10^−4^, and batch size 8. The Adam optimizer and mean squared error loss were used during training. To deal with overfitting, dropout between layers and early stopping were used. Test performance was assessed using mean absolute error to compare results for different modalities and combinations of modalities.

## 4. Results

### 4.1 Results for Experiment 1

For chronological age prediction, we ran the models for epoch 50, 75 and 100 twice. The average of MAEs from both runs was calculated to compare the modalities. The common trend in the results showed that the best performance for each modality was around epoch 75. As we wanted to see the effect of training the model for more epochs, we kept the patience of early stopping at 100. The increase in the value of MAE after epoch 75 indicates that early stopping can be utilized based on validation loss to prevent overfitting at that point.

T1w derived BrainAge had the best performance (MAE: 5.474) on our test dataset at 75 epochs. FA maps gave the best MAE of 3.19 for 50 epochs, whereas MD and RD maps gave the best MAE of 4.31 and 3.99 respectively at 75 epochs. AD maps had the best performance at 75 epochs with a MAE of 3.69.

Based on the above results, we calculated the delta brain age gap between predicted age and actual age for both CN/MCI and CN/AD. Using the features actual age, sex and diagnosis, we used linear regression to predict the brain age gap. The prediction of this gap is better for CN vs AD in comparison to CN vs MCI, as expected.

### 4.2 Results for Experiment 2

For the 2D slice encoder network, the performance was similar to the 3D CNN architecture for T1w scans, relative to the DWI metrics. The best MAE was in the case of DWI-RD, whereas the worst was in the case of DWI-FA.

### 4.3 Results for Experiment 3

For chronological age prediction using both modalities merged, the performance of the model was worse as compared to running on a single modality; the main reason may be a limited number of training data when the scans are stacked together. The performance of the concatenated model was better than the stacked data model.

### 4.4 Results for Experiment 4

For the different clinical scores, the best prediction performance was for MMSE, with an MAE of 1.54 for DWI-RD scans. The worst performance was for ADAS13, with an MAE of 5.01 for DWI-FA scans. For individual modalities, the best MAEs were for MMSE, and the performance was similar for ADAS11 and ADAS13 prediction.

## 5. Conclusions

For BrainAge Prediction, diffusion-weighted images may offer better results for deep learning algorithms using 3D CNNs, compared to CNNs trained on T1w images alone. For 2D slice networks, the results are slightly better for T1w scans, but still comparable. For chronological age prediction using fused data modalities, the performance of the model was worse than training on a single modality; this was surprising, but the main reason may be higher dimension of the input data, relative to the limited amount of training examples, when the scans are stacked together. The 3D CNN architecture can also be used to predict clinical scores such as ADAS11, ADAS13 and MMSE from both T1w and DWI images, with better results coming from the DWIs. The experiments show that DWI image modalities can be used as a close substitute for T1w images for various prediction tasks. In future we plan to include more matched T1w and DWI data, from larger datasets such as the UK Biobank [26], and more diverse datasets such as ENIGMA [27], to further evaluate model performance. Future work will also assess the added value of imaging protocols such as DWI TDF (a scalar measure computable from a tensor distribution function fitted to single shell diffusion MRI), for improving model performance. We also plan to test data harmonization approaches based on generative adversarial networks and related approaches, which can make data from different scanners or protocols more comparable - hence helping create a holdout dataset for testing model performance.

## Acknowledgments

This research was supported in part by the NIH under grants: RF1 AG057892, P41EB015922, R01AG058854, U01AG068057, U01AG024904 (to ADNI), and the U.S. Department of Defense. We thank the ADNI investigators and their public and private funders for creating and publicly disseminating the ADNI dataset.

